# Chemi-Proteome Language Attention Network Empowers Fragment-Based Ligand Interactome and Binding Sites Discovery with Evidence

**DOI:** 10.64898/2026.08.26.747036

**Authors:** Bin Liao, Jixiao He, Mingzhu Zhao, Xiaozhen Cui, Yunxia Cui, Chuanqiao Dong, Hongyan Sun, Liang Zhang, Jian Zhang

## Abstract

Deep learning has accelerated drug discovery, yet most existing models are trained using in vitro affinity datasets and consequently remain disconnected from the cellular context in which functional ligand–protein interactions occur. This limitation hinders the ability to reflect the complexity of native interactomes and characterize biological responses to molecular perturbation. Here we introduce C-PLANK (Chemi-Proteome Language Attention NetworK), a deep learning framework trained on fragment–protein interactions profiled directly in living cells using fully functionalized fragment (FFF) chemoproteomics. C-PLANK combines physicochemical embeddings with a bilinear attention network (BAN) to model both global cellular context and local residue–atom interactions, generating interpretable interaction fingerprints. Particularly, C-PLANK incorporates Cellular Interaction State Index (CISI), a systems-level evidential metric that contextualizes the biological plausibility of each predicted interaction against the global cellular interaction landscape. Across 431 ligand interactomes curated from eight independent chemoproteomic studies, C-PLANK consistently outperformed current state-of-the-art interaction prediction frameworks under both random and cold-protein evaluation settings. The inferred interaction fingerprints aligned with orthogonal evidence from structure-based pocket predictions, co-crystal structures, and cellular binding-site annotations. C-PLANK further generalized to unseen ligands. In a cellular target-focused discovery campaign, C-PLANK identified a previously unrecognized ligand that was subsequently advanced into an active chemical probe acting as a SIRT3 agonist in cellular assays. By learning directly from cellular chemoproteomics, C-PLANK moves beyond isolated interaction prediction toward cellular interaction-state modelling, establishing a computational foundation for future digital-twin frameworks in drug discovery.

## Introduction

Deep learning has transformed drug discovery by reducing reliance on labor-intensive experimental screening and enabling the exploration of vast chemical spaces through data-driven prediction models^1-5^. Supported by the rapid accumulation of drug–target interaction (DTI) datasets generated from high-throughput screening (HTS), machine learning frameworks have demonstrated impressive efficiency in identifying ligand-protein interactions and prioritizing candidate compounds for downstream development^4,6^. As a result, significant efforts have focused on developing computational models capable of accurately predicting molecular interactions and functional effects.

Current DTI prediction methods differ primarily in how proteins and ligands are represented. Structure-based approaches leverage experimentally determined or computationally predicted three-dimensional structures paired with molecular descriptors of ligands^7^. However, the incomplete coverage of high-resolution structures across the proteome limits the applicability of these methods^8^. Sequence-based approaches address this challenge by representing proteins as amino-acid sequences and ligands as simplified molecular-input line-entry system (SMILES) strings, with convolutional neural networks (CNNs) learning local sequence and chemical patterns^9-13^. Other methods adopt graph representations of proteins and compounds and employ graph neural networks (GNNs) to capture topological relationships^14-17^. More recently, transformer and attention-based architectures have demonstrated strong performance by modeling long-range dependencies and cross-modal interactions between proteins and ligands^18-20^. Collectively, these advances have established deep learning as a powerful paradigm for molecular interaction prediction and drug discovery.

Despite these sucesses, existing models are mostly trained on in vitro affinity datasets, such as BindingDB^21^, Davis^22^ and BioSNAP^23^, that describe interactions between compounds and isolated proteins under controlled biochemical conditions (Supplementary Information, Section 1). While valuable, these datasets provide only a partial view of biological reality. They do not capture the dynamic cellular environment in which ligand-protein interactions naturally occur, including contextual effects arising from protein complexes, post-translational regulation, and signaling activity. Furthermore, many affinity datasets lack experimentally verified negative interactions, requiring computational generation of negative samples that may introduce false negatives and skew model training^24^. Consequently, modern AI systems effectively learn pairwise interaction patterns but remain relatively disconnected from the biological systems that are the ultimate goal of modeling. Emerging concepts such as digital twins emphasize the need of computational representations that integrate heterogeneous data streams, quantify uncertainty, and characterize cellular states under perturbation. In drug discovery, such representations should ideally inform how compounds interact with the proteome in cellular contexts, rather than focusing on pair-wised binding events. Achieving this goal requires novel modeling exploiting data that directly reflects biological interactions at the systems level.

Fragment-based ligand discovery (FBLD) provides a promising route toward such system-aware modeling^25^. Unlike HTS, which screens large collections of fully elaborated compounds, FBLD interrogates the cellular proteome using chemically simple fragments (∼150–300 Da) that efficiently sample chemical space^26-30^. FBLD libraries often yield higher hit rates and uncover ligandable sites on proteins that are difficult to access using conventional screening approaches^30, 31^. However, fragment–protein interactions are typically weak and transient, making them challenging to detect withy traditional biochemical assays^30-32^. The development of fully functionalized fragment (FFF) probes has transformed this landscape. By incorporating photo-reactive and bioorthogonal handles, FFF probes enable covalent capture of transient interactions and proteome-wide profiling of ligand binding in living cells through chemoproteomics^25^. As a result, FFF datasets provide a unique view of context-dependent and reversible interactions within the cellular proteome (Fig. 1a). Nevertheless, mass spectrometry (MS)–based chemoproteomics remains resource-restricted, and the synthesis and deployment of diverse libraries are costly^33-38^. Computational models capable of learning from cellular chemoproteomic perturbations therefore have the potential to substantially reduce experimental burden while accelerating fragment-based discovery^30, 32,39^.

**Fig. 1.**
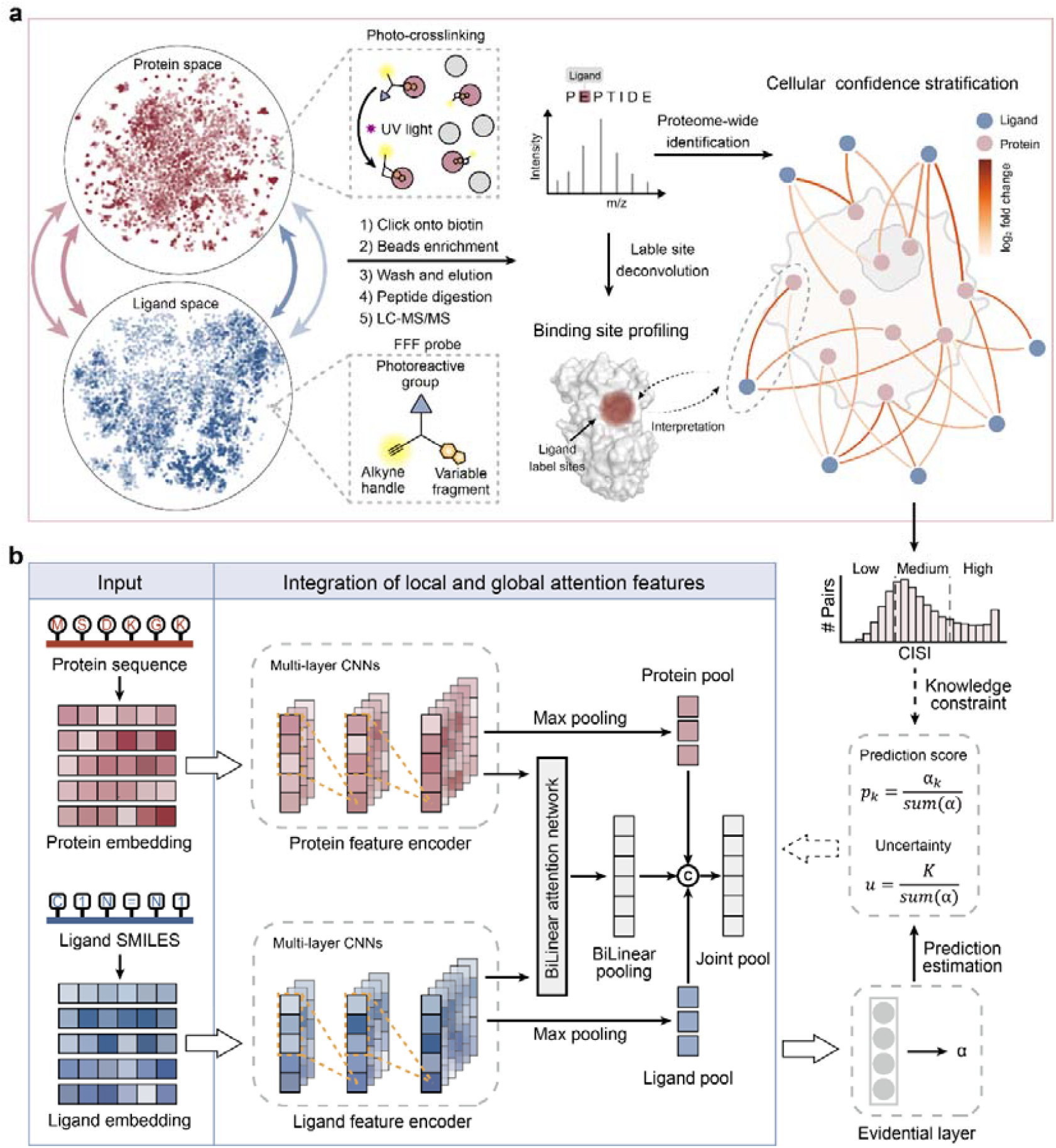
An overview of C-PLANK for learning fragment-protein interactions from chemoproteomic data. **a**, Schematic of FFF-based ligand discovery strategy and its integration with deep learning. In FBLD, FFF probes derived from ligand space consist of a small molecule library to inquire ligandable proteins from protein space. Each FFF probe contains a variable fragment for reversible reactions with proteins, a photoreactive group active upon UV irradiation, and an alkyne handle attached to a biotin tag by click chemistry for enriching labeled proteins. Coupled with LC-MS/MS analysis, a proteome-wide fragment–protein interactome and ligandable binding sites can be achieved. **b**, Architecture of C-PLANK. Protein and ligand embedding are derived from protein sequence and ligand SMILES respectively. The abstract features of protein and ligand are respectively extracted by a 1D-CNN block and fused via BAN module to generate local attention, which is concatenated with global attention to obtain a parameter α in evidential layer. Prediction score and uncertainty are estimated by α and constrained by chemoproteomics prior knowledge as well.

Here we introduce Chemi-Proteome Language Attention NetworK (C-PLANK), a deep learning framework designed to learn directly from cellular chemoproteomic datasets generated by FFF probes. Unlike conventional DTI models trained on biochemical affinity measurements, C-PLANK is built to model proteome-wide interaction landscapes of ligands within cellular context. The framework combines protein language representations, physicochemical features, and a bilinear attention mechanism to capture both global contextual signals and local residue-atom interactions. Moreover, we integrate evidential deep learning (EDL)^11, 40^ with Cellular Interaction State Index (CISI), a system-level evidential metric that contextualizes compound-protein interactions against the cellular interaction landscape. C-PLANK moves beyond conventional pairwise DTI prediction toward a computational framework for cellular interaction-state modelling, establishing a foundation for future digital-twin approaches to drug discovery.

## Result

### Architecture of the C-PLANK framework

C-PLANK was developed as a framework that predicts and interprets fragment–protein interactions within the context of the cellular proteome. Unlike conventional drug–target interaction (DTI) models that learn pairwise relationships from in vitro affinity measurements, C-PLANK learns directly from chemoproteomic perturbation data generated in living cells, allowing it to model interactions within cellular interaction landscape (Fig. 1a and Supplementary Information, Section 2). To achieve this, the framework comprises three major components: ligand and protein encoders for feature extraction, a bilinear attention network (BAN) decoder for interaction-state learning, and an evidential layer for uncertainty-aware inference of cellular interactome (Fig. 1b).

For a given ligand–protein pair, C-PLANK first transforms protein sequences and ligand SMILES into chemically informed representations before integrating them through attention-based interaction learning. Protein sequences are represented using protein language model ESM-2 with curated physicochemical descriptors (Supplementary Fig. 1a). This dual representation captures both evolutionary context and intrinsic biochemical properties related to ligand recognition. In parallel, ligand fragments are encoded at the atom level through a learnable embedding matrix augmented with atom-specific physicochemical features (Supplementary Fig. 1b), preserving fragment-level chemical diversity and functional-group information. These protein and ligand representations are independently processed by stacked one-dimensional convolutional neural networks (1D-CNNs) to extract intra-sequence and intra-molecular patterns (Fig. 1b). Each encoder generates two complementary outputs: (i) a max-pooled global representation that summarizes long-range contextual information, and (ii) a position-resolved feature matrix that retains residue- and atom-level information for subsequent interaction-state modelling (Fig. 1b). Through this design, C-PLANK simultaneously captures global properties of proteins and ligands, and the fine-grained features required to describe specific interaction events.

To model local interaction states, C-PLANK employs a bilinear attention network (BAN) that learns pairwise relationships between protein residues and ligand atoms (Supplementary Fig. 1c). The BAN module generates an interaction attention map highlighting residue–atom hotspots together with a pooled local interaction vector. Whereas the global representations characterize the overall biochemical context, the BAN-derived features encode localized interaction evidence. The combination of these two components yields a joint representation that integrates system-level context with residue-level interaction determinants, enabling the prediction of fragment–protein interaction propensities while simultaneously producing interpretable interaction fingerprints.

Beyond interaction prediction, C-PLANK explicitly estimates the confidence and uncertainty associated with the interaction state within the cellular interactome. To contextualize these estimates within the cellular chemoproteomic landscape, we developed a Cellular Interaction State Index (CISI). CISI is a systems-aware evidential metric defined by the enrichment of a fragment–protein interaction relative to the global promiscuity of the corresponding ligand and protein across the chemoproteomic dataset (Fig. 1a-b). Rather than reflecting model confidence alone, CISI quantifies how strongly interaction is supported by the observed cellular interaction network. This yields positive and negative interactions that are stratified into high-, medium-, and low-CISI groups (Supplementary Fig. 2a-b; Supplementary Information, Section 3). These interaction-state priors were integrated into the evidential framework that outputs the evidence parameter α, enabling C-PLANK to jointly learn interaction propensity, uncertainty, and biological credibility (Fig. 1b). Consequently, the model produces not only a prediction of whether an interaction is likely to occur, but also an estimate of how consistently that interaction is supported by the broader cellular interaction landscape.

**Fig. 2.**
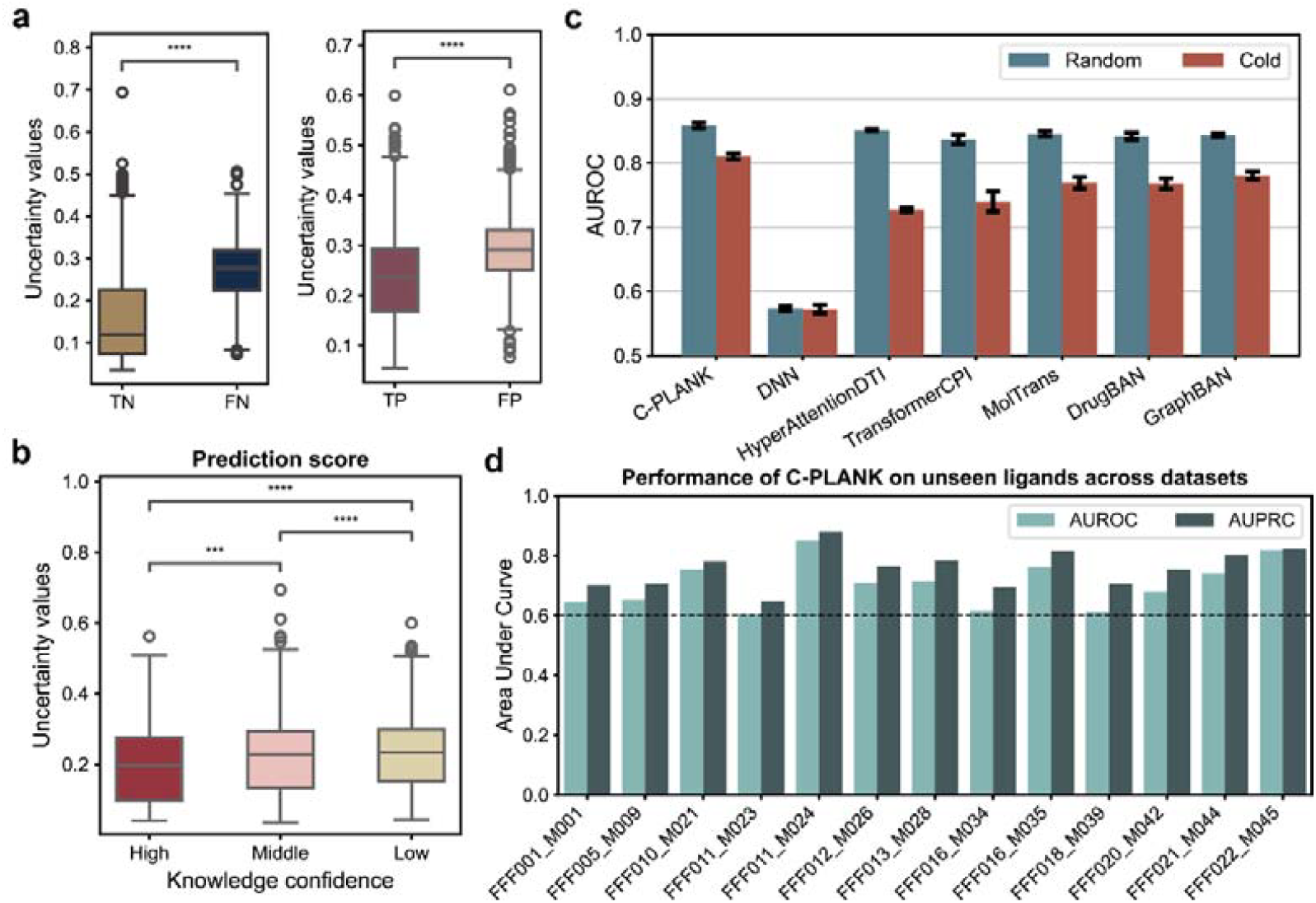
Performance evaluation and comparison on custom dataset. **a**, Distribution of uncertainty values in predicted negative (left) and positive (right) samples. **b**, Distribution of uncertainty values under different confidence score groups according to prior knowledge. **c**, AUROC result across baselines in random and cold split (10-fold cross validation). **d**, Bar plot showing AUROC and AUPRC values of 13 unseen ligands with specific methods. Only paired ligand name (front) and method id (back) with AUROC > 0.6 and AUPRC > 0.6 are displayed. The vertical bars and black lines denote as mean values and standard deviation respectively. TN: true negative; FN: false negative; TP: true positive; FP: false positive. For a and b, Wilcoxon rank-sum test is leveraged to measure statistical significance, ***P < 1e-3 and ****P < 1e-4.

### Chemoproteomic dataset curation and evaluation strategies

To train C-PLANK with a dataset that captures cellular interaction states, we curated chemoproteomic studies employing fully functionalized fragment (FFF) probes together with inactive negative-control probes^35^. We prioritized fragment-inactive controls over competitor- or enantiomer-based controls because they enable more stringent discrimination between genuine fragment-dependent enrichment and nonspecific pulldown events. In addition, we focused on probes containing diazirine photoreactive groups, the most widely adopted photoreactive chemistry in FFF-based chemoproteomics, to maximize consistency across experimental studies and reduce systematic biases arising from probe architecture.

In total, we collected 431 ligand interactomes from eight independent chemoproteomic studies (Supplementary Data 1), derived from Wozniak et al.^39^ and Chem(Pro)2^41^. Unlike conventional DTI datasets, where each interaction is treated as an isolated observation, these datasets provide proteome-scale measurements of cellular responses to chemical perturbations, thereby defining a global interaction landscape. To mitigate bias introduced by highly recurrent ligands or proteins, we constructed a balanced training dataset comprising 90,494 interactions between 407 ligands and 2,629 proteins, based on the frequency distribution of ligands and proteins across the interactomes (Details provided in Supplementary Information, Section 2)^35^. The final dataset was randomly split into training, validation, and test sets at an 8:1:1 ratio. To rigorously evaluate whether models learn transferable biological representations rather than memorizing target identities, we employed two complementary evaluation schemes. The first was a standard random split, which measures predictive performance under in-distribution conditions. The second was a stringent cold-protein split, where proteins were clustered according to 3-mer frequency profiles and assigned to mutually exclusive training, validation, and test sets (Supplementary Information, Section 4). This setting evaluates the ability of models to infer interaction states for previously unseen proteins, better reflecting real-world ligand discovery scenarios.

Model performance was primarily assessed using the area under the receiver operating characteristic curve (AUROC) and the area under the precision-recall curve (AUPRC). Accuracy, precision, and recall were additionally reported for the optimal model. To improve statistical robustness, we performed 10-fold cross-validation, generating independent training, validation, and test partitions for each fold. Model selection was based on the highest validation AUROC, and corresponding performance metrics were subsequently calculated on the held-out test set. Importantly, the curated chemoproteomic resource serves not only as a training benchmark but also as a quantitative representation of the underlying cellular interaction landscape, forming the basis for uncertainty-aware inference in C-PLANK.

### C-PLANK integrates cellular interactions to assess confidence and uncertainty

A defining feature of C-PLANK is its ability to assess the confidence and uncertainty of predicted interactions using the cellular interactome landscape. We first examined the relationship between predictive correctness and model uncertainty. Predictions were categorized into four groups according to the confusion matrix: true negative (TN), false negative (FN), true positive (TP) and false positive (FP) (Fig. 2a). Both TP and TN predictions exhibited significantly lower uncertainty than FP and FN predictions, indicating that the evidential framework successfully distinguishes reliable from unreliable interaction-state predictions. This result demonstrates that uncertainty estimates generated by C-PLANK are informative rather than merely reflecting model confidence.

We next investigated whether uncertainty estimates were consistent with CISI derived from chemoproteomic prior knowledge. Interaction pairs were stratified into high-, medium-, and low-CISI groups, representing different levels of support from the observed chemoproteomic landscape. We found a clear monotonic relationship between CISI and uncertainty, with high-CISI interactions consistently exhibiting lower uncertainty than medium- and low-CISI interactions (Fig. 2b). This observation suggests that interactions strongly supported by proteome-wide chemoproteomic evidence are also assigned greater certainty by C-PLANK. Thus, the chemoproteomic prior contributes information that cannot be recovered from prediction outcomes alone.

Collectively, these findings indicate that CISI provides an orthogonal source of biological evidence for uncertainty quantification. Rather than relying exclusively on model outputs, C-PLANK calibrates interaction-state estimates using information derived from the global cellular interaction landscape, resulting in uncertainty measures that are both interpretable and biologically grounded.

### Benchmark performance, ablation study, and generalization of C-PLANK

Next, we benchmarked C-PLANK against seven baselines, including a basic deep neural network (DNN) and six state-of-the-art models: HyperAttentionDTI^12^, TransformerCPI^20^, MolTrans^19^, DrugBAN^42^ and GraphBAN^43^ (details are provided in Supplementary Information, section 6). Across random splits, C-PLANK consistently achieved the best overall performance, yielding superior AUROC and accuracy while maintaining competitive precision and recall (Table 1 and Fig. 2c). In contrast, the shallow DNN performed substantially worse, highlighting the importance of deep contextual representations and explicit interaction modelling for cellular states inference. Under the more challenging cold-protein split, performance decreased for all methods, reflecting the difficulty of transferring interaction knowledge to previously unseen proteins. Nevertheless, C-PLANK maintained its lead, outperforming the strongest baseline by approximately 1–2% across AUROC and precision (Table 1; Fig. 2c). These improvements indicate that C-PLANK learns more transferable representations of fragment-induced cellular interaction states and generalizes more effectively across the proteome. Collectively, these results demonstrate that C-PLANK not only improves interaction prediction accuracy but also generates robust and uncertainty-aware representations of the cellular interaction landscape.

**Table 1.** Comparison with baseline models on custom dataset with random or cold split.

|  | Method | Accuracy (sd) | Precision (sd) | Recall (sd) | AUROC (sd) | AUPRC (sd) |
| --- | --- | --- | --- | --- | --- | --- |
| Random split | DNN* | 0.5620 ± 0.0026 | 0.5897 ± 0.0091 | 0.4084 ± 0.0101 | 0.5732 ± 0.0036 | 0.5790 ± 0.0069 |
| | HyperAttentionDTI* | <u>0.7638 ± 0.0014</u> | **0.7565 ± 0.0010** | 0.7779 ± 0.0026 | <u>0.8515 ± 0.0010</u> | <u>0.8359 ± 0.0015</u> |
| | TransformerCPI* | 0.7500 ± 0.0074 | 0.7200 ± 0.0106 | 0.8191 ± 0.0087 | 0.8367 ± 0.0068 | 0.8205 ± 0.0093 |
| | MolTrans* | 0.7594 ± 0.0029 | 0.7203 ± 0.0092 | 0.8490 ± 0.0200 | 0.8454 ± 0.0040 | 0.8279 ± 0.0061 |
| | DrugBAN* | 0.7511 ± 0.0084 | 0.6895 ± 0.0146 | <u>0.9148 ± 0.0156</u> | 0.8415 ± 0.0057 | 0.8213 ± 0.0097 |
| | GraphBAN* | 0.7505 ± 0.0076 | 0.6848 ± 0.0123 | **0.9293 ± 0.0123** | 0.8437 ± 0.0028 | 0.8261 ± 0.0050 |
| | C-PLANK | **0.7718 ± 0.0047** | <u>0.7424 ± 0.0088</u> | 0.8328 ± 0.0156 | **0.8586 ± 0.0042** | **0.8413 ± 0.0055** |
| Cold split | DNN* | 0.5609 ± 0.0069 | 0.5855 ± 0.0147 | 0.4142 ± 0.0164 | 0.5723 ± 0.0062 | 0.5763 ± 0.0109 |
| | HyperAttentionDTI* | 0.6286 ± 0.0032 | 0.6719 ± 0.0049 | 0.4949 ± 0.0076 | 0.7270 ± 0.0028 | 0.6700 ± 0.0039 |
| | TransformerCPI* | 0.6839 ± 0.0133 | <u>0.6729 ± 0.0229</u> | 0.7225 ± 0.0623 | 0.7511 ± 0.0219 | 0.6991 ± 0.0249 |
| | MolTrans* | 0.7076 ± 0.0094 | 0.6696 ± 0.0089 | 0.8197 ± 0.0476 | 0.7690 ± 0.0089 | 0.7157 ± 0.0124 |
| | DrugBAN* | 0.7051 ± 0.0084 | 0.6439 ± 0.0103 | **0.9174 ± 0.0157** | 0.7678 ± 0.0083 | 0.7148 ± 0.0131 |
| | GraphBAN* | <u>0.7158 ± 0.0060</u> | 0.6506 ± 0.0075 | <u>0.9137 ± 0.0103</u> | <u>0.7806 ± 0.0062</u> | <u>0.7340 ± 0.0075</u> |
| | C-PLANK | **0.7272 ± 0.0048** | **0.7100 ± 0.0207** | 0.7730 ± 0.0672 | **0.8106 ± 0.0042** | **0.7788 ± 0.0089** |

The matrices are presented as mean ± standard deviation (std). The best model under each metric is marked bold while the secondary best model is underlined. \*Represents the baseline framework has statistically significant difference with C-PLANK on AUROC, which is measured by $p$ -value ( $p < 0.05$ ) using a two-sided paired t-test.

To assess the contributions of individual architectural components, we constructed six C□PLANK variants by selectively removing physicochemical descriptors, chemoproteomic prior knowledge, and global or local attention mechanisms. For the one-sided attention variants, we replaced the bilinear attention network (BAN) with a Transformer-style decoder analogous to TransformerCPI, in which cross-attention is used to fuse ligand and protein embeddings and infer interaction features^20^. This design enabled us to evaluate whether the performance gains of C□PLANK arise from its bilinear interaction modelling or from attention mechanisms more generally.

Across all evaluation settings, the complete C□PLANK architecture consistently outperformed every ablated variant (Table 2). The largest performance decreases were observed when either the global contextual representation or the local BAN-derived interaction representation was removed, indicating that accurate modelling of cellular interaction states requires both system-level context and fine-grained residue–atom interaction evidence. Removing physicochemical descriptors from either proteins or ligands consistently impaired model performance, demonstrating that C□PLANK effectively leverages intrinsic biochemical properties as complementary signals to learned representations. In addition, replacing BAN with Transformer-style cross-attention significantly reduced performance. This observation highlights the ability of bilinear attention to capture sparse and localized residue–atom interaction patterns that characterize fragment-based chemoproteomic datasets. Such interactions are often weak and transient, making their detection particularly dependent on fine-grained modelling of pairwise interaction evidence.

**Table 2.**
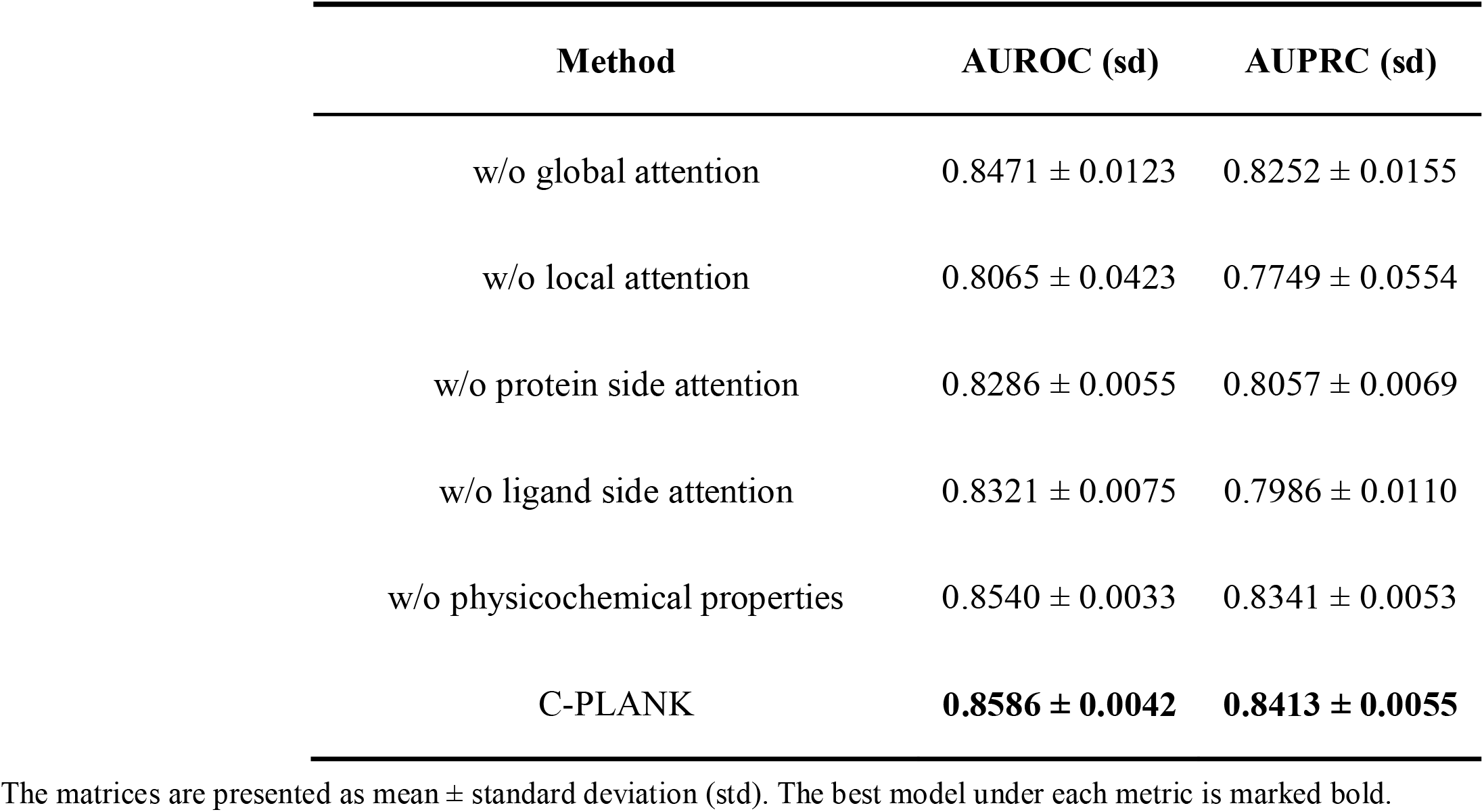
The ablation study in AUROC and AUPRC on custom dataset.

To evaluate the generalization of C-PLANK, we exploited independent chemoproteomic datasets containing 24 previously unseen ligands, many of which also involved unseen proteins (Supplementary Data 1). To avoid evaluation bias, balanced test sets were generated by sampling negative proteins from those detected but not significantly enriched relative to matched negative-control probes. Under moderate performance criteria (AUROC and AUPRC ≥ 0.6), C□PLANK successfully reconstructed the interaction landscapes of 13 of 24 ligands (Fig. 2d), demonstrating substantial transferability across distinct chemoproteomic studies. All experiment details and prediction performance are available in Supplementary Information section 7. Notably, the interactome generated by ligand FFF011 (M024) achieved an AUROC of 0.8497, indicating that the learned interaction-state representations can remain informative to truly independent experiments.

### Interpretability analysis of C-PLANK

Through its bilinear attention network (BAN), C□PLANK generates atom- and residue-level attention scores that identify the molecular determinants contributing to a predicted interaction state (Fig. 1b). To evaluate whether these inferred interaction fingerprints correspond to genuine biological signals rather than model artifacts, we compared them against multiple orthogonal experimental and computational sources of evidence.

We first leveraged Dizco, chemoproteomic platform that provides proteome-wide binding-site mapping of FFF probes in living cells^39^. The seven Dizco ligands used in our generalization analysis (Supplementary Information, section 7) provided an opportunity to systematically examine how C□PLANK distributes attention across fragment architectures. Across all ligands, C□PLANK consistently assigned the highest interaction-state importance to the variable fragment moiety, followed by the diazirine photoreactive group, while the alkyne enrichment handle received substantially lower attention scores (Fig. 3a). This hierarchy closely mirrors the chemistry of FFF probes: the fragment moiety mediates initial reversible target engagement, whereas the diazirine group becomes activated upon UV irradiation and covalently captures nearby residues to stabilize transient interactions (Fig. 1a). The low attention assigned to the alkyne handle further suggests that C□PLANK focuses on chemically functional regions relevant to biological activity.

**Fig. 3.**
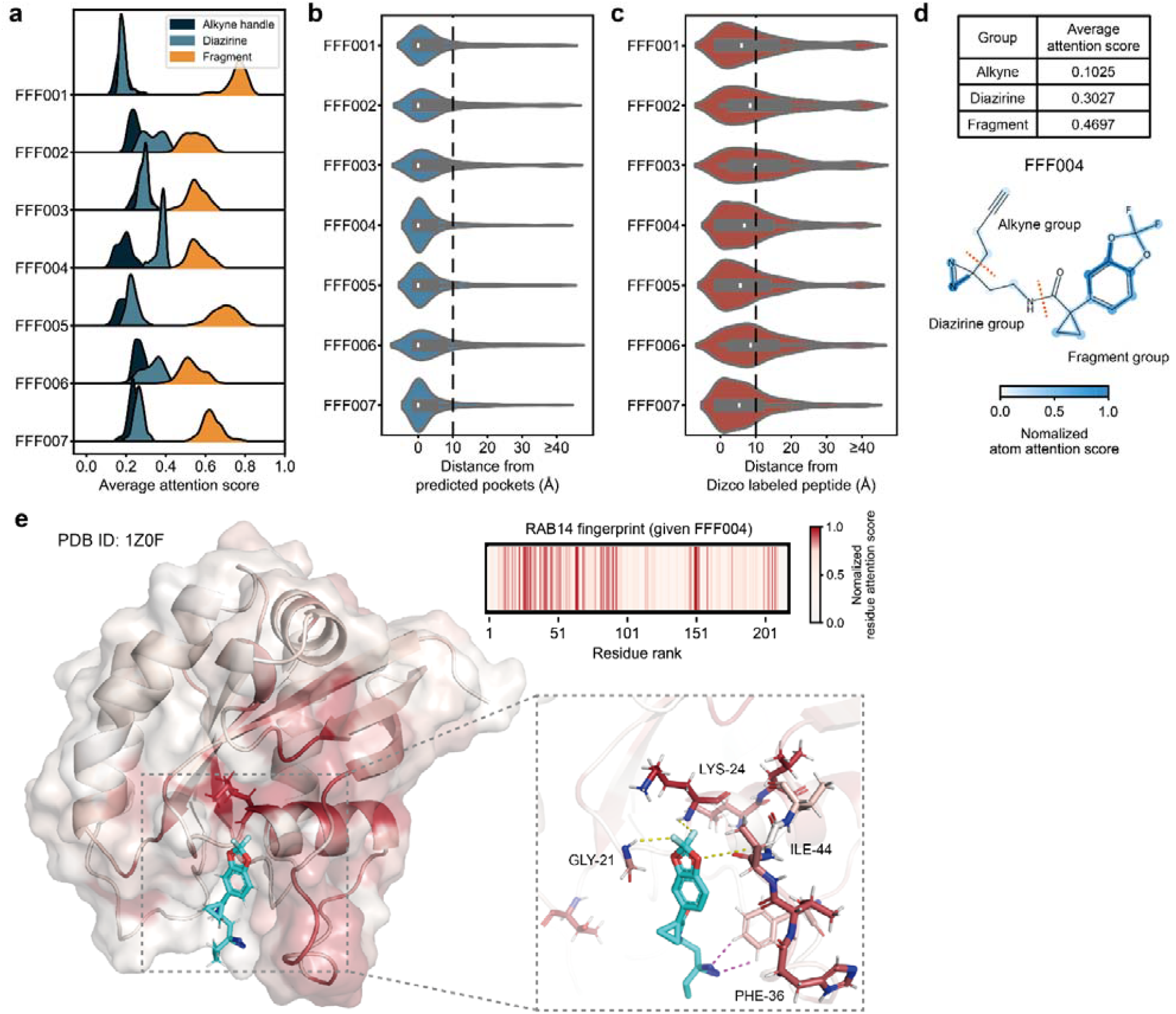
Interpretable fingerprint provides reveal interacting patterns of ligand and protein. **a**, Average attention score distribution of alkyne handle, diazirine and variable fragment group across seven ligands. **b-c**, Distribution of distance between highlighted residues of C-PLANK and high-confidence predicted pockets from Fpocket **(b)** and Dizco labeled peptides **(c). d**, Normalized atom attention scores from C-PLANK for FFF004, grouped by chemical moiety (alkyne handle, diazirine photoreactive group and variable fragment). Scores are projected onto the FFF004 chemical structure (bottom). **e**, Residue attention scores projected onto the RAB14 structure (PDB ID: 1Z0F^47^) in complex with FFF004 (docking-based pose). Inset plot (top right) shows residue-level attention fingerprint across RAB14 sequence. Bottom inset plot displays zoomed view of the GDP-binding pocket region, highlighting key interacting residues (K24, G21, I44, F36) with FFF004. For protein mapping in **e**, the residues with attention scores in top 20% are highlighted with dark red while the remaining regions are colored with gradient light red.

To systematically evaluate residue-level interpretability, we projected C□PLANK attention scores onto experimentally determined protein structures from the Protein Data Bank (PDB)^44^ or AlphaFold predictions^7^ and quantified their spatial relationship to Dizco-labeled binding sites (Supplementary Information, Section 8). After min–max normalization of attention values, residues with high attention scores (>0.8) were compared with experimentally identified labeling sites. We compared C□PLANK attention maps against ligand-binding pockets predicted by Fpocket^45^. To ensure high confidence, only pockets with druggability scores greater than 0.5 were retained (Supplementary Information, Section 9). High-attention residues identified by C□PLANK showed substantial overlap with residues constituting these predicted pockets (Fig. 3b), despite the model relying solely on sequence-derived representations rather than explicit structural coordinates. This observation indicates that the learned interaction-state representations preserve information associated with protein ligandability. A substantial fraction of high-attention residues directly overlapped with Dizco-labeled positions (0–3 Å), while the majority were located within 10 Å of the experimentally mapped sites (Fig. 3c). Given the finite labeling radius of diazirine-based photoreactive probes^46^, this level of spatial agreement strongly suggests that the inferred interaction states capture biologically meaningful binding determinants. Taken together, these analyses demonstrate that C□PLANK learns biologically meaningful interaction-state representations that are supported by independent chemoproteomic evidence, structural pocket predictions, and functional annotations.

To further illustrate the interpretability of C-PLANK outputs, we investigated two representative systems with co-crystal structures and experimental binding-site support. In the first labeling event between FFF004 and RAB14 (PDB ID: 1Z0F^47^), C-PLANK assigned the highest average attention scores to variable fragment group of FFF004, followed by diazirine photoreactive group (Fig. 3d). Notably, the 2,2-difluoro-1,3-benzodioxole moiety within fragment group received the highest attention scores, indicating its critical contribution to FFF004-RAB14 interaction. On the protein side, we utilized molecular docking to inquire possible binding poses between FFF004 and RAB14, referencing the known position of GDP-binding pocket in 1Z0F^47^. The results revealed favorable interactions of 2,2-difluoro-1,3-benzodioxole with K24, I44 and G21 residues, which are also emphasized by C-PLANK (Fig. 3e). In addition, C-PLANK assigned a high attention score to F36 proximal to diazirine, which was independently confirmed by Dizco.

### Application of C-PLANK discovers a novel SIRT3 agonist

To evaluate whether cellular interaction-state modelling can support prospective ligand discovery, we applied C□PLANK to identify modulators of SIRT3, a mitochondrial NAD^+^-dependent deacetylase that is localized to the mitochondrial matrix^48^. Through regulation of mitochondrial protein acetylation, SIRT3 contributes to diverse biological processes, including aging, neurodegeneration, mitophagy, and metabolic homeostasis^49^. Despite its therapeutic potential, few selective SIRT3 agonists have been reported^50, 51^, highlighting the need for efficient discovery strategies.

We utilized C□PLANK to perform a screening campaign against the large fragment library described above (Fig. 4a). Ligands were first ranked according to their predicted interaction-state scores toward SIRT3, and the top 50 candidates (approximately the top 1% of the screened library) were retained for further evaluation. To prioritize compounds with favorable properties, we incorporated additional criteria reflecting both confidence and developability. Specifically, candidates were filtered according to predictive uncertainty (<0.25), fragment promiscuity (<40%) estimated using a previously reported machine-learning framework^35^, and calculated Walden–Crippen LogP values (<2.5). This multi-criteria prioritization strategy integrates interaction propensity, uncertainty, and chemoproteomic context, providing a practical example of how cellular interaction-state modelling guides decision making in ligand discovery.

**Fig. 4.**
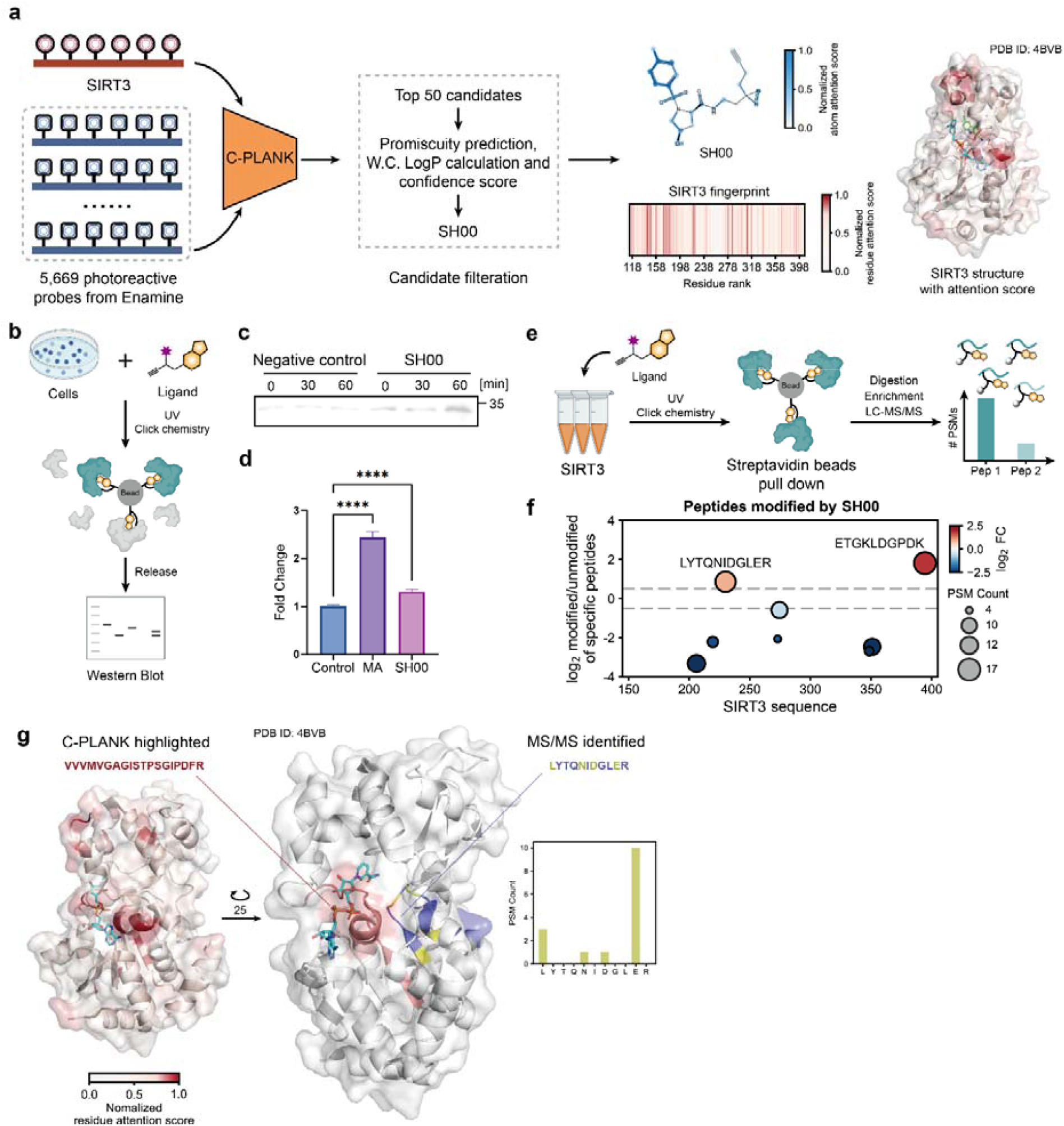
Discovery of a SIRT3 agonist using C-PLANK. **a**, Workflow of SIRT3 screening for a candidate modulator discovery and SH00 was eventually identified as an effective candidate. **b**, Schematic of validating interaction between SIRT3 and SH00 in cells via western blot. **c**, Gel-based FBLD analysis of Hela cells treated with negative control or SH00 under gradient incubation time (0, 30, 60 min with concentration of 20 μM). **d**, Fold change in SIRT3 deacetylase activity induced by the positive control 1-methylbenzylamino amiodarone (MA) and SH00 relative to the negative control probe, as measured by the fluorescence detection (FD) assay. **e**, Quantitative proteomics for validating the binding event and mapping the binding sites. **f**, Bubble heatmap of SIRT3 peptides, showing the log_2_ intensity FC of specific peptides with versus without SH00 modification. The color denotes log_2_ intensity FC and the dot size denotes number of PSMs of modified peptides. **g**, Matching the binding regions predicted by C-PLANK and identified by quantitative proteomics (PDB ID: 4BVB^52^). Visualized interpretability of C-PLANK reveals the binding sites with high normalized residue scores (left). Middle plot displays C-PLANK highlighted peptide (red) and chemoproteomics identified labeled peptide in which labeled residues are colored with yellow while the remainders are colored with purple. The left plot shows PSM count of labeled residues for significant peptide related to **f**. Data presented are mean ± SD, n = 3 biological replicates for **c** with one-way Anova analysis, ***P < 1e^-5^.

Following this filtering process, SH00 emerged as a relatively promising candidate, exhibiting low predicted promiscuity, favorable physicochemical properties, and a high interaction-state score for SIRT3 (Fig. 4a; Supplementary Fig. 3a; Supplementary Data 2). To experimentally validate the prediction, we performed chemoproteomic profiling in HEK293T cells using SH00 (Fig. 4b). Relative to negative control probe, SH00 produced significant enrichment of SIRT3, confirming its binding capability (Fig. 4c). Furthermore, fluorescence detection (FD) assays demonstrated that SH00 enhanced the deacetylase activity of SIRT3, indicating agonistic activity rather than simply target engagement (Fig. 4d). These results validate the practical utility of C□PLANK for identifying biologically active modulators from large chemical libraries.

Analysis of the corresponding interaction fingerprint revealed that C□PLANK assigned the highest atom-level attention scores to the variable fragment moiety, with the benzenesulfonyl group being particularly emphasized (Fig. 4a), suggesting that this substructure contributes substantially to the inferred SIRT3 interaction state. In parallel, we performed chemoproteomic analysis to map the labeling regions of SH00 on SIRT3 (Fig. 4e). The peptide-spectrum match (PSM) results indicate reproducible SH00 labeling events on SIRT3 (Fig. 4f and Supplementary Fig. 3b). To compare these observations with the inferred interaction state by C-PLANK, residue attention scores were projected onto the SIRT3 structure. C□PLANK highlighted residues including G145, G147, T150, and P151, which are located proximal to the NAD^+^-binding region (Fig. 4g). Notably, the experimentally labeled peptides identified by chemoproteomics localized within the photolabeling radius of these highlighted residues (Fig. 4g), establishing strong spatial agreement between computational inference and experimental measurements.

Although the exact labeling positions did not perfectly overlap with the highest-attention residues, such offsets are expected given both the labeling radius of photoaffinity probes and the tendency of C□PLANK to preferentially emphasize interaction-relevant fragment moieties rather than capture residues themselves^35, 36, 46^. Importantly, the inferred interaction fingerprint correctly localized the interaction state to the functionally relevant region surrounding the NAD^+^ binding pocket. This convergence between prediction, chemoproteomic validation, and structural context provides evidence that C□PLANK captures biologically meaningful determinants of target engagement. In this case study, C□PLANK translates cellular interaction-state representations to enable the discovery of a previously unrecognized SIRT3 binder with agonistic activity.

### Structural optimization and functional assessment of SIRT3 agonists

Given the weak SIRT3 activation observed for SH00SIRT3 (1.3-fold at 100 μM, Fig. 4d), systematic structural optimization was undertaken (Fig. 5a, Supplementary Fig. 3c and Supplementary Information, section 10 and 11). Removal of the photo-crosslinking moiety and simplification of the stereochemical scaffold yielded SH01, while subsequent phenyl-ring incorporation afforded SH02. Both analogues exhibited enhanced SIRT3 activation relative to the parent hit SH00. Next, a bioisosteric replacement strategy was applied to optimize the amide moiety of the parent scaffold. Among the resulting analogues, SH04, featuring a 1,2,4-oxadiazole ring, showed the most pronounced activity, inducing a 3.94-fold activation of SIRT3 at 100 μM. SIRT3 Encouraged by this result, a series of para-substituted phenyl analogues bearing electron-donating and electron-withdrawing substituents were prepared to probe the SAR around the oxadiazole moiety. Most derivatives showed markedly enhanced activity at 10 μM, with the para-chloro analogue SH06 displaying the highest potency and inducing a 2.1-fold activation of SIRT3.

**Fig. 5.**
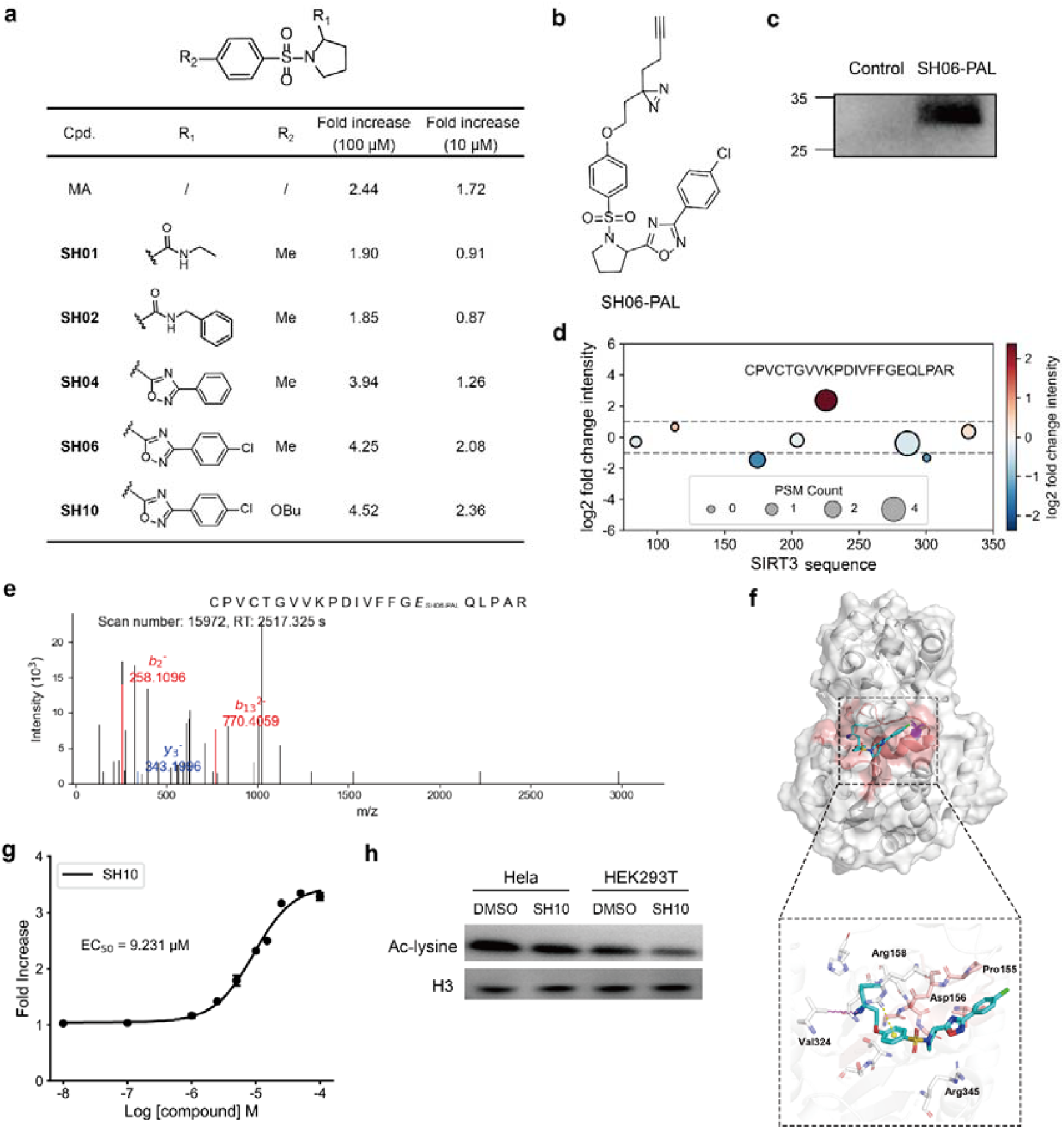
Optimization of SH00 enhances the activation ability toward SIRT3. **a**, Chemical structures of SH00 and the development to SH10. The bottom row shows SIRT3 activity FC of each probe. **b**, Chemical structure of SH06-PAL. **c**, Gel-based FBLD analysis of Hela cells treated with SH06-PAL and negative control probe (20 μM). **d**, Bubble heatmap of SIRT3 peptides, indicating the significant peptide modified by SH06-PAL, analogous to **Fig. 4f. e**. MS/MS spectra example for peptide of CPVCTGVVKPDIVFFGEQLPAR with SH06-PAL modification on 17E identified by FragPipe. **f**, Molecular docking of SH06-PAL and SIRT3. The key residues (red) are highlighted that surround SH06-PAL (cyan) (PDB ID: 4JSR^53^). The purple region represents labeled residue identified by FragPipe. **g**, Concentration–response analysis of SH10-mediated SIRT3 activation. A non-linear fit function is applied. **h**, Immunoblot analysis of DMSO and SH10 in Hela and HEK 293T cell lines, indicating the deacetylation level of SHO10 (10 μM, 48 h). Data presented are mean ± SD, n = 3 biological replicates for **g**. Significant peptides in **d** denotes log_2_ FC > 0.5.

Subsequently, we introduced a photoreactive group equipped with an alkyne handle into SH06 to generate SH06-PAL for profiling its binding site on SIRT3 (Fig. 5b). We first demonstrated that SH06-PAL efficiently enriched SIRT3 from cell lysates (Fig. 5c) and subsequently applied a chemoproteomic workflow to identify the labeled peptide and the corresponding binding residue, E233 (Fig. 5d, e). Molecular docking was then performed to elucidate the binding mode of SH06-PAL, revealing a binding conformation in which the interacting residues were near the labeled peptide (Fig. 5f). Finally, substitution of the methyl group on the sulfonyl phenyl ring with a butoxy group yielded SH10 (Supplementary Fig. 3d), resulting in further improvement in SIRT3 activation at both 100 and 10 μM. Notably, SH10 retained detectable activity at 1 μM, producing a 1.1-fold activation of SIRT3 and an EC50 value of 9.23 μM (Fig. 5g). Surface plasmon resonance (SPR) analysis further validated the direct binding of SH10 to SIRT3, yielding a Kd value of 10.9 μM (Supplementary Fig. 3e). Consistent with its biochemical activity, SH10 enhanced SIRT3-mediated deacetylation in both HeLa and HEK-293T cells, as evidenced by a marked reduction in global lysine acetylation levels (Fig. 5h). To the best of our knowledge, SH10 exhibits a favorable activity profile compared with previously reported SIRT3 agonists, with a single digit micromolar EC□□ and a relatively high Emax.

## Discussion

Chemoproteomics enables comprehensive profiling of fragment-protein interactions across proteome in negative biological systems. The expanding scale of data from fragment-based screening necessitates robust deep learning tools to accelerate exploration of fragment space. To this end, we proposed C-PLANK, a deep learning framework that learns global features from the physicochemical properties of ligands and proteins while employing BAN to capture local interaction patterns, thereby comprehensively characterizing fragment-protein interactions. Furthermore, the interpretability of C-PLANK advances our understanding of fragment-protein interactions and provides critical guidance for fragment optimization. C-PLANK has demonstrated reliable generalizations to unseen ligands. In a large-scale ligand screening for SIRT3, C-PLANK identified promising modulator which has been further optimized and developed into a potent SIRT3 agonist, SH10, underscoring its potential in ligand discovery.

Despite these advancements, C-PLANK has some limitations warranting further refinement. Currently, our training dataset encompasses interactome from only 407 ligands, limiting the diversity of interaction patterns that can be learned. Casually assembling fragment-protein interaction data from disparate sources for training could expand the scope of non-specific or indirectly enriched proteins, introducing data distribution shifts that compromise model performance. To address this, future efforts could focus on developing automated chemoproteomics benchmarking methods to enhance identification throughput and speed and yield standardized datasets. Additionally, developing advanced statistical methods could mitigate batch effects arising from differences in instruments and experimental conditions, facilitating seamless integration of multi-source data.

This version of C-PLANK relies on 1D protein sequences and ligand SMILES as inputs, primarily due to the limited availability of reliable high-resolution protein structures. Although C-PLANK has impressively demonstrated its effectiveness in elucidating fragment-protein interactions using 1D sequence data, incorporating high-quality 3D structural information of proteins could capture additional spatial information, augmenting its performance and interpretability. This is particularly promising given the advancements in protein structure prediction models such as AlphaFold^7^ and RoseTTAFold^54^.

Furthermore, C-PLANK presently processes isolated ligand–protein pairs. However, recent research suggests that targets pulled down by FFF probes are interconnected, often as part of protein complexes^36^. Therefore, future improvements could extend C-PLANK to accept multiple proteins as inputs, integrating protein-protein interaction network information to improve model performance and enable cross-domain data fusion.

In conclusion, by learning from cellular data and producing mechanistically grounded outputs, C□PLANK moves toward a future in which AI models can reason not only about molecular interactions but also about the negative biological systems in which those interactions occur.

## Method

### C-PLANK architecture

#### Initialization of protein and ligand representations

For given 1D protein sequences with different lengths of amino acids, C-PLANK sets a maximum length *N*_*p*_ of protein sequences, with longer sequences truncated and shorter sequences padded with zeros to maintain a consistent input length for batch training. Each protein contains two types of features at residues: pretrained protein language features from ESM-2 and physicochemical properties. The pretrained features derived from the last layer embedding of ESM-2 are represented as 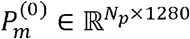 and the physicochemical properties, including molecular weight, polarity, hydropathicity, pKa, and topology, are encoded into 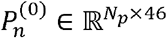. A protein feature encoder is implemented to map both residue features to a latent embedding space to compute protein feature embedding 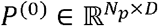.

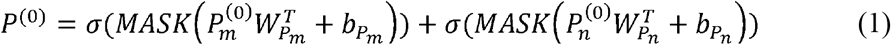

where *D* is the feature hidden embedding dimension in C-PLANK, 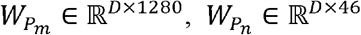 and 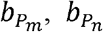, are the learnable weight matrices and bias vectors respectively. The *MASK* denotes an operation of masking the padding position to reduce the interference and is *σ* a non-linear activation function.

Given the input of ligand SMILES containing different number of atoms, C-PLANK also sets a fixed length *N*_*l*_ of ligand atoms for batch training. Considering the light molecular weight of FFF probes, *N*_*l*_ can be a moderate number which would be enough to cover all FFF probes in library and reduce padding length. The ligand feature embedding can be described by 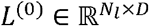:

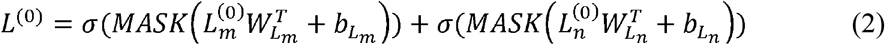

where 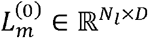 represents an atom-type features derived from a learnable embedding matrix consisting of core atoms of FFF probes (C, N, O, F, P, S, Cl, Br and I) and the other minor atoms, and 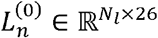 denotes the physicochemical properties matrices of atoms, including LogP, charges, topological polarity surface area (TPSA) and hybridization. 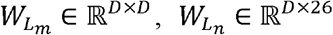 and 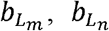, are the learnable weight matrices and bias vectors respectively. Here, *MASK* not only excludes the influence of paddings but also distinguishes the token of paddings and minor atoms, which share same label in 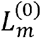.

#### CNN block for proteins and ligands

In this module, C-PLANK implements two independent 1D-CNN blocks, both comprising three consecutive convolution layers, to extract the local patterns of *P*^(0)^ and *L*^(0)^ from latent space respectively. With the increasing convolution kernel size *K*, enlarged receptive fields enable this approach capturing more abstract relationships of overlapping *K*-mer amino acids from short range to long range for protein and identifying more diverse features of photoactive, alkyne handle and fragment group for ligand. The CNN architecture can be described as:

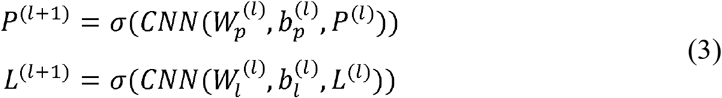

where 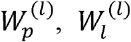 and 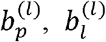 are the learnable weight matrices and bias vector respectively. The *P*^(*l*)^ and *L*^(*l*)^ are the representation matrices in *l*th layer of protein and ligand respectively. The output matrices of 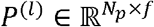 and 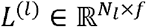 are both subsequently processed into two types of representations, where *f* is the output dimension of 3-layer CNN. On the one hand, a global-max pooling module is leveraged to convert *P*^(*l*)^ and *L*^(*l*)^ into vectorized features *v*_*p*_ and *v*_*l*_ along amino acids and atoms dimensions respectively, which are concatenated as global attention embedding Ψ_*global*_ ℝ ∈^2*f*^ for prediction. On the other hand, in order to separate feature extractor by CNN block and pairwise attention module, C-PLANK transforms *P*^(*l*)^ and *L*^(*l*)^ into global attention matrices 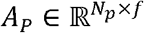 and 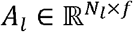 respectively as follows:

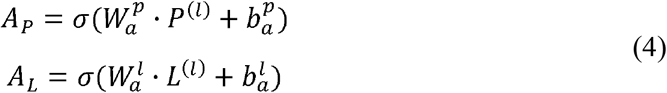

where 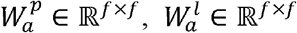 and 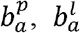 are the learnable weight matrices and bias vector respectively.

#### Bilinear attention network for pairwise interaction learning

BAN was originally proposed to address the visual question answering (VQA) problems, which involved aligning multimodal input like visual and natural language question information, and providing an answer toward the graph-text matching^55^. Recently, BAN has been extended to the prediction filed of DTI and demonstrated its major benefits^42, 43^. In C-PLANK, BAN is introduced to model the pairwise interactions between protein amino acids and ligand atoms, thereby enhancing the capture of local interaction features. Notably, in C-PLANK, the BAN module enables accepting the unimodal sequential hidden embeddings of protein and ligand as input, since BAN excels at learning fine-grained joint representations between any pair of feature modalities, regardless of their original form. The BAN module is composed of two core components: the bilinear attention map, which integrates the hidden representations of ligand and protein to generate a pairwise attention-weight matrix, and the bilinear pooling, which applies the attention-weight matrix to extract the joint local feature representation of ligand and protein.

The first element of BAN module is bilinear attention map which learns the pairwise interactions and creates a single head attention-weight matrix 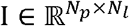 from attention matrices 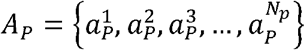 and 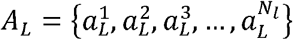, in which 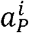 and 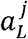 represent the *i*th and *j*th substructural attention embedding of a given protein and ligand respectively. Specifically, each element I_*ij*_ in I is the low-rank output of pairwise local interaction between *i*th element of protein and *j*th element of ligand, computed as:

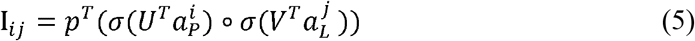

where *p* ∈ ℝ^*K*^ denotes a learnable *K*-dimention interaction weight vector, *U* ∈ ℝ ^*f*×*K*^ and *V* ∈ ℝ ^*f*×*K*^ are the learnable weight matrices for protein and ligand representations respectively and ° stands for Hadamard product, i.e., the element-wise product. The bilinear attention map can be extended to multiple heads to comprehensively learn the pairwise interaction details by defining various learnable *p*_*g*_ weight vector, where *g* is the index of heads, and sharing *U* and *V* across heads, which eventually leads to a multiple attention-weight matrix 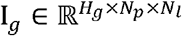, where *H*_*g*_ is the number of heads.

In the second element of BAN module, a bilinear pooling is introduced to generate joint representation of local interaction, *f* ∈ ℝ^*K*^, in which the *k*th component can be computed as:

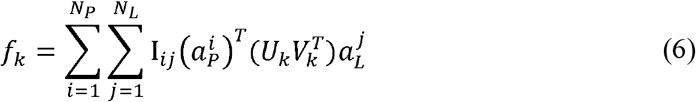

where *U*_*k*_ and *V*_*k*_ denote the *k*th column of *U* and *V* respectively. In multi-head module, the final joint representation Ψ_*local*_ ∈ ℝ^*K*/*s*^ represents as follows:

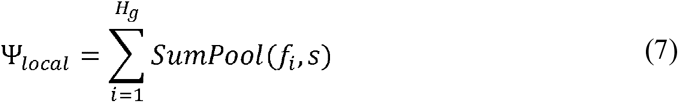

where *i* is the *i*th head, *SumPool* denotes an operation of sum pooling on the *i*th output of joint representation *f*_*i*_ with a stride of *s*, which compresses the dimension of joint representation from *K* to *K*/*s*. For balance with Ψ_*global*_, the local attention vector Ψ_*local*_ is configured such that *K*/*s* = 2*f*.

#### Interpretation layer

The comprehensive integrated bilinear attention map I_*g*_ is leveraged to calculate the attention scores for each protein residue and ligand atom. In the first step, I_*g*_ undergoes average pooling over the dimension of ligand and protein to generate local attention matrices 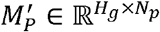 and 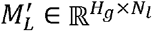 respectively. The second step considers the weighted attribution of various heads on local attention and computes the integrated local attention matrices 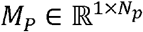 and 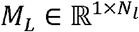 as:

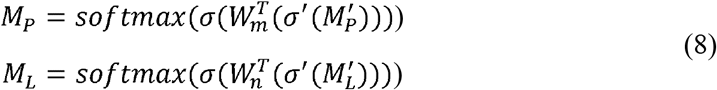

where 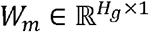 and 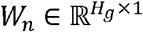 are learnable SoftMax weight matrices used to compute the weighted sum over the features of protein residues and ligand atoms respectively, across attention heads. The outermost SoftMax layer considers the actual length of protein and ligand while the remainder is masked with zero.

#### Evidential layer and objective function

For computing the interaction probability, we integrate global and local interaction vectors and feed them into an evidential layer which consists of a multi-layer fully connected neural network (FCNN), eventually obtaining evidence *e*:

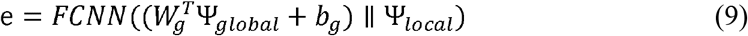

where *W*_*g*_ ∈ ℝ^*f*×*f*^ denotes a learnable weight matrix which transfers global attention vector into a more abstract representation, *b*_*g*_ is a bias vector and ‖ represents a concatenation operation on global and local attention vectors.

The evidence *e* meets requirement of non-negative via softplus transformation and corresponds to a Dirichlet distribution with parameter *α*_*k*_ = *e*_*k*_ + 1, where *k* denotes *k*th class. The evidence *e* and uncertainty *u* follow the relationship:

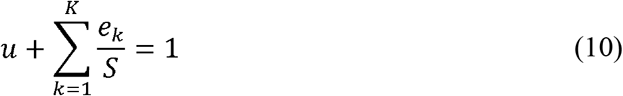

where 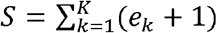 is normalization coefficient. Therefore, equation (10) can be rewritten as:

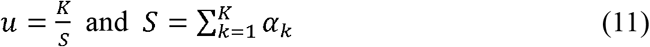

The prediction probability 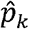 of *k*th class represents as the mean of corresponding Dirichlet distribution:

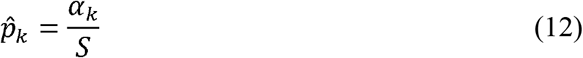

The entire learnable parameters are jointly optimized with the objective of minimizing the loss consisting of EDL loss ℒ_*E*_ and knowledge loss ℒ_*K*_:

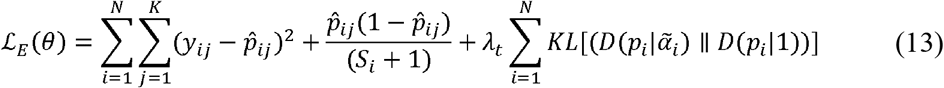

where *y*_*ij*_ is the ground truth label of *i*th fragment-protein interaction in *j*th class, *λ*_*t*_ denotes an anneal hyperparameter relevant to training epoch 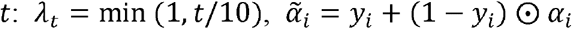 is the Dirichlet parameters keeping the evidence of *y*_*i*_ unchanged and pushing Dirichlet distributions 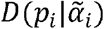 toward a uniform distribution *D*(*p*_*i*_|1) which represents a completely uncertain situation.

To constrain the uncertainty for each interaction *i* using prior knowledge, we defined knowledge loss ℒ_*K*_ according to groups with different CISI.

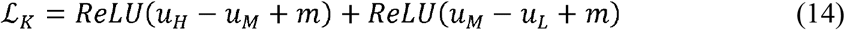

where *u*_*H*_, *u*_*M*_ and *u*_*L*_ are the mean uncertainty values of high-, medium- and low-CISI groups respectively. Eventually, the total loss function ℒ = ℒ_*E*_ + *λ*_*t*_ ℒ_*K*_ .

### Experimental setting

#### Training data

To train C-PLANK, we constructed a custom dataset comprising 407 photoreactive ligands profiled proteome-wide. The positive fragment-protein pairs met the following criteria: (1) the log_2_ fold change (ligand/control) > 2.3 (fivefold enrichment); (2) the *p*-value < 0.05; (3) mdfClass ≥ 2; (4) The number of unique peptides ≥ 2 and (5) the enriched proteins have been annotated in Uniprot. Ultimately, this yielded 45,247 positive fragment–protein pairs involving 332 ligands and 2,629 proteins. To prevent overfitting and biased performance evaluation, we selected an equal number of negative pairs for each protein by matching negative ligands according to their promiscuity, resulting in balanced positive and negative classes. The final dataset contained 90,494 fragment-protein pairs and was randomly split with the ratio of 0.8/0.1/0.1 for training, validation and test dataset respectively.

#### Hyperparameters of C-PLANK

Hyperparameters of C-PLANK was tuned using Optuna 4.5.0^56^ package with Tree-structured Parzen Estimator (TPE) sampler. Consequently, we selected AdamW as the optimizer with the learning rate of 1 × 10^-4^ and weight decay of 0.01. A CosineAnnealingLR scheduler dynamically adjusted the learning rate across training epochs, with a maximum of 100 epochs permitted for C-PLANK. Protein sequences and ligand SMILES were truncated or padded to maximum lengths of 1,200 residues and 60 atoms, respectively, with both protein and ligand embeddings set to a feature dimension of 64. The 1D-CNN block consists of three layers with increasing channel dimensions of [32, 64, 128] and kernel sizes of [3, 7, 11], enabling the model to progressively capture increasingly larger receptive fields. The BAN module employed 4 attention heads and an output dimension of 256 to capture fine-grained local interaction details. The optimal model checkpoint of C-PLANK was selected at the epoch with the best AUROC (Area Under Receiver Operating Characteristic) score on validation set, which would be used to evaluate model performance on test set. The hyperparameter configurations are provided in Supplementary Information, section 5.

### Experiments and assays for SIRT3 agonist discovery

#### Chemoproteomics experiment methods

Hela cells were treated with probes of 0/10/20 μM in serum-free media for 60□min followed by UV of 312□nm irradiation for 20□min at 4□°C to stabilize probe-protein interaction. The prepared reaction regent was added to samples prior click chemistry, including 30□μL tris[(1-benzyl-1H-1,2,3-triazol-4-yl)methyl]amine consisting of 1.7□mM TBTA in DMSO-tBuOH (1:4 v/v), 10□μL CuSO4 of 50□mM, 5□μL biotin enrichment tag of 10□mM and 10□μL tris(2-carboxyethyl)phosphine (TCEP) of 50□mM. The click reaction was carried out for 1□h at room temperature with a gentle shake. Following addition of 2□mL methanol:chloroform (4:1) and 1□mL DPBS to precipitate proteins, pelleted proteins were washed twice with 2□mL methanol:chloroform before being resuspended in 500□μL 6□M urea in DPBS. Protein samples were added with 130□μL 10% SDS and diluted to 5.5□mL with DPBS. Prewashed 100□μL streptavidin agarose beads were added to each sample, and proteins were enriched for 1.5□h at room temperature with gentle rotation. Streptavidin beads were pelleted by centrifugation and washed once with 5□mL 0.2% SDS in DPBS, twice with DPBS and twice with H_2_O. Washed beads were transferred to tubes using 100□mM triethylammonium bicarbonate (TEAB) buffer and 200□μL of a trypsin digest solution (2□μg trypsin, 1□mM CaCl_2_ in 100□mM TEAB) was added to each sample. Proteins were allowed to digest overnight at 37□°C with a gentle shake. Labeled peptides were released from streptavidin beads with 2□×□1□h incubations in 200□μL 2% formic acid followed by 1□h incubation in 400□μL 1% formic acid in 50% acetonitrile.

#### LC-MS/MS analysis

Nanoflow capillary columns (150□µm I.D.) with pulled nanoESI emitters were packed to 150□mm at high pressures with 2□µm diameter, 100□Å pore size C18 particles. Samples were analyzed with a Vanquish Neo UHPLC (Thermo Scientific) coupled to an Orbitrap Astral mass spectrometer (Thermo Scientific) using a NanoSpray Flex source (Thermo Scientific). A source voltage of 1900 V was used for all experiments. Mobile phase A and B were 0.1% formic acid in water (Fisher Scientific, Optima LC-MS grade) and 0.1% formic acid/80% acetonitrile (Merck, Optima LC-MS grade), respectively. The column was heated to 50□°C. The gradient was initiated at 4% B with a flow rate of 2.5 μL/min. Solvent B was increased to 5% at 0.5 min and to 8.5% at 0.9 min while the flow rate was reduced to 0.8 μL/min. Solvent B was then increased from 8.5% to 25% over 13.0 min, to 35% over 6.9 min, and to 55% over 0.4 min. Solvent B was subsequently ramped to 99% over 0.5 min while the flow rate was increased to 2.5 μL/min, followed by a 0.9 min column wash at 99% B. The system pressure was maintained below 1000 bar throughout the analysis. For initial DDA experiments on the Orbitrap Astral MS, MS1 spectra were collected in the Orbitrap every 0.6□s at a resolving power of 240,000 over m/z 350–1350 with a normalized AGC target of 500% (Absolute AGC Value: 5e6) and a maximum injection time of 10□ms. The MIPS filter was applied with Peptide mode and “Relax Restrictions when too few Precursors are Found” set to True. Precursors were filtered to charges states 2-6. A Dynamic Exclusion filter was applied with 10□s duration and 10ppm low and high mass tolerance and exclude isotopes set to True. An intensity filter was applied with a minimum precursor intensity of 5000 required for selection. MS2 scans were collected in the Astral mass analyzer with an isolation window of 1.6□m/z, normalized collision energy of 28, a first mass of 120 m/z, an AGC target of 500% (Absolute AGC Value: 5e4), and a maximum injection time of 3□ms. MS raw data were searched against SIRT family (downloaded from Uniprot, 2024.09.12) with FragPipe platform (v24.0). The offset search workflow was applied to inquiry any amino acids labeled with negative control (+ 294.206), SH00 (+ 519.252) and SH06-PAL (+ 640.233). The delta mass information can be checked in assigned modifications column. The other parameters were set as default.

#### SIRT3 protein expression and purification

Recombinant human SIRT3 (residues 118–399) bearing an N-terminal His tag was expressed in Escherichia coli using a pET-28a expression vector. Protein expression and purification were performed as previously described^57, 58^. Briefly, protein expression was induced with 0.5 mM isopropyl β-D-1-thiogalactopyranoside (IPTG; I5502, Sigma-Aldrich, USA). After cell harvest, the lysate was clarified by centrifugation, and the recombinant protein was purified by Ni–NTA affinity chromatography.

#### Fluorescence detection (FD) assay

SIRT3 deacetylase activity was measured using a fluorescence detection (FD) assay as previously described1. Briefly, 2μM SIRT3 was incubated with 10 μM fluorogenic substrate peptide Z-(Ac)Lys-AMC and 1 mM NAD□ (V900401, Sigma-Aldrich, USA) in the absence or presence of the indicated compounds, with the final DMSO concentration maintained at 1%. After incubation at 37 °C for 30 min, the reaction was terminated by the addition of trypsin (10 μM; T4799, Sigma-Aldrich, USA) and nicotinamide (2 mM; 72340, Sigma-Aldrich, USA). Fluorescence intensity was measured using a Synergy Neo microplate reader (BioTek, USA) with excitation and emission wavelengths of 360 and 460 nm, respectively.

#### Surface Plasmon Resonance (SPR) Assay

The binding affinity of **SH10** for SIRT3 was determined by surface plasmon resonance (SPR) using a Biacore 8K instrument (Cytiva, USA) equipped with a CM5 sensor chip (BR-1005-30, Cytiva). Recombinant SIRT3 was diluted in 10 mM sodium acetate buffer (pH 5.0) and immobilized onto the sensor surface by standard amine coupling using an Amine Coupling Kit (BR-1000-50, Cytiva). **SH10** was prepared in running buffer containing 5% DMSO at final concentrations of 1.5625, 3.125, 6.25, 12.5, 25, 50, and 100 μM. Binding analyses were performed at 25 °C with a flow rate of 30 μL min□^1^. The association and dissociation phases were monitored for 180 and 250 s, respectively. Sensorgrams were processed using Biacore 8K Manager software (Cytiva), and kinetic parameters, including the equilibrium dissociation constant (K_D), were obtained by global fitting of the concentration-dependent binding data to a 1:1 Langmuir binding model.

## Data availability

All training and evaluation data used in C-PLANK can be found at https://github.com/Bin-cc/C-PLANK. The The information of commercial fully functionalize compounds is available at https://enamine.net/compound-collections/photoaffinity-compounds. The protein crystal structures used in C-PLANK can be found at https://www.rcsb.org/ by searching their PDB IDs. The large-scale chemoproteomics fragment-protein interactions information are provided in https://www.science.org/doi/abs/10.1126/science.adk5864, Tables S1-2. The Dizco-labeled peptides and functional sites utilized for distance comparison are available at https://www.nature.com/articles/s41589-023-01514-z, Supplementary Data 1. The reset ligand information can be searched in Chem(Pro)2: https://idrblab.org/chemprosquare by entering corresponding ligand names.

## Code availability

The implementation code and details of C-PLANK are freely available at https://github.com/Bin-cc/C-PLANK, along with the environment configurations and installation instructions for running the code.

## Acknowledgements

This work was supported by grants from Innovative Drug Research and Development National Science and Technology Major Project (2025ZD1803104 to JZ), Shanghai Municipal Health Commission (2025ZHYL038 to JZ), Lingang Laboratory (LG8888 to JZ), Shanghai Action Plan for Science, Technology and Innovation Field of Computational Biology (24JS2830100 to JZ), the Research Grants Council of Hong Kong (11101025, 11102322, 11104422, C1024-22GF, T12-101/23-N, C1041-24EF to LZ), a grant from Innovation and Technology Fund of Hong Kong (ITS/169/23 to LZ), a grant from The Tung Foundation Biomedical Sciences Centre (9609314 to LZ).

## Author contributions

B.L., L.Z. and J.Z. conceived and designed the study. B.L., J.H. and M.Z. conducted the experiments and analyzed the data. X.C, Y.C., C.D., and H.S. provided technical support. B.L. and J.H. prepared the figures and drafted the manuscript. L.Z. and J.Z edited and revised the manuscript.

